# Dynamic Grouping of Ongoing Activity in V1 Hypercolumns

**DOI:** 10.1101/2024.12.20.629848

**Authors:** Rui Zhang, Jiayu Wang, Xingya Cai, Rendong Tang, Haidong D. Lu

**Affiliations:** State Key Laboratory of Cognitive Neuroscience and Learning, IDG/McGovern Institute for Brain Research, Beijing Normal University, Beijing, China; Department of Neurology, Zhongshan Hospital, Institute for Translational Brain Research, Fudan University, Shanghai, China

**Keywords:** macaque, V1, V2, spontaneous activity, neuron ensemble, functional map, two-photon calcium imaging, functional connectivity

## Abstract

Neurons’ spontaneous activity provides rich information about the brain. A single neuron’s activity has close relationships with the local network. In order to understand such relationships, we studied the spontaneous activity of thousands of neurons in macaque V1 and V2 with two-photon calcium imaging. In V1, the ongoing activity was dominated by global fluctuations in which the activity of majority of neurons were correlated. Neurons’ activity also relied on their relative locations within the local functional architectures, including ocular dominance, orientation, and color maps. Neurons with similar preferences dynamically grouped into co-activating ensembles and exhibited spatial patterns resembling the local functional maps. Different ensembles had different strengths and frequencies. This observation was consistent across all hypercolumn-sized V1 locations we examined. In V2, different imaging sites had different orientation and color features. However, the spontaneous activity in the sampled regions also correlated with the underlying functional architectures. These results indicate that functional architectures play an essential role in influencing neurons’ spontaneous activity, and can be explained by a network model that integrates diverse horizontal connections among similar functional domains.

## Introduction

When we examine a single neuron, its spontaneous activity appears to be random. However, when more neurons are examined simultaneously, we found that neurons’ spontaneous activity often correlate with each other, especially for neurons close-by (Leopold et al. 2003; Smith & Kohn 2008; Smith & Sommer 2013). The correlation strength varies with neurons’ selectivity features and their locations (Maier et al. 2010; Okun et al 2015). Such variation may due to the intrinsic cortical functional networks, like orientation maps (Arieli et al. 1995, 1996; Tsodyks et al. 1999; Singh et al. 2008; Cai et al. 2023). However, direct evidence for such dependency is still lacking. For non-human primates, there are many functional architectures in their sensory cortices. For example, there are orientation, color, and ocular dominance maps in area V1. In the hypercolumn model Hubel and Wiesel proposed, a column under a 1-mm^2^-sized surface contains full cycles for all these features (Hubel & Wiese 1974). How do these nested networks affect neuron’s spontaneous activity? Whether a single neuron’s activity is determined by the neuron’s spatial location within these functional maps? The answers to these questions remain unknown.

Two-photon calcium imaging provides a way to image hundreds to thousands of neurons within a mm^2^-sized cortical region. It has been used in studying spontaneous activity in different animal models, for example in mice (Goltstein et al. 2015; Mizuno et al. 2018), rats (Ch’ng & Reid 2010), cats (Ch’ng & Reid 2010), and ferrets (Smith et al. 2018). To our knowledge, there is no report of two-photon study of spontaneous activity in non-human primates. Previously, we used two-photon calcium imaging and studied neurons’ visual response features (Tang et al. 2020). Here, we examined spontaneous activity with the same technique. We characterized neurons’ spontaneous activity with respect to their spatial context in the local functional maps, as well as their selectivity to visual features. As a classical model for brain research, macaques have been studied on spontaneous activity in both micro (electrophysiology) and macro (fMRI) scales. Two-photon imaging has the unique resolution and scale to bridge these findings.

## Results

In 3 macaques, we injected AAV virus to express GCaMP6s proteins in multiple locations in V1 and V2. A chronic optical chamber with a diameter of 13-mm was implanted (Figure 1A&B). In the subsequent anesthetized experiments, we imaged calcium signals with a 16X objective. The frame size was 0.83 x 0.83 mm, collected at 1.3Hz frame rate. Imaging depth was 180-270 μm. In each experiment, imaging of spontaneous activity always started first, normally lasted for 70 min (∼5000 frames), when the animal’s eyes were closed. After that, visual responses of the same group of neurons were collected. Visual stimuli were drifting gratings of different colors and orientations, presented monocularly or binocularly. Each stimulus was presented for 2 seconds. About 400-600 cells (average 569 cells) can be identified from each imaging site (e.g., Figure 1C). Neuron’s fluorescent activity time courses were filtered (0.005-0.3Hz), transformed to dF/F, and z-scored. Following results were all based on these n x m (cell number x frame number) matrices (e.g., Figure 1F).

**Figure 1.**
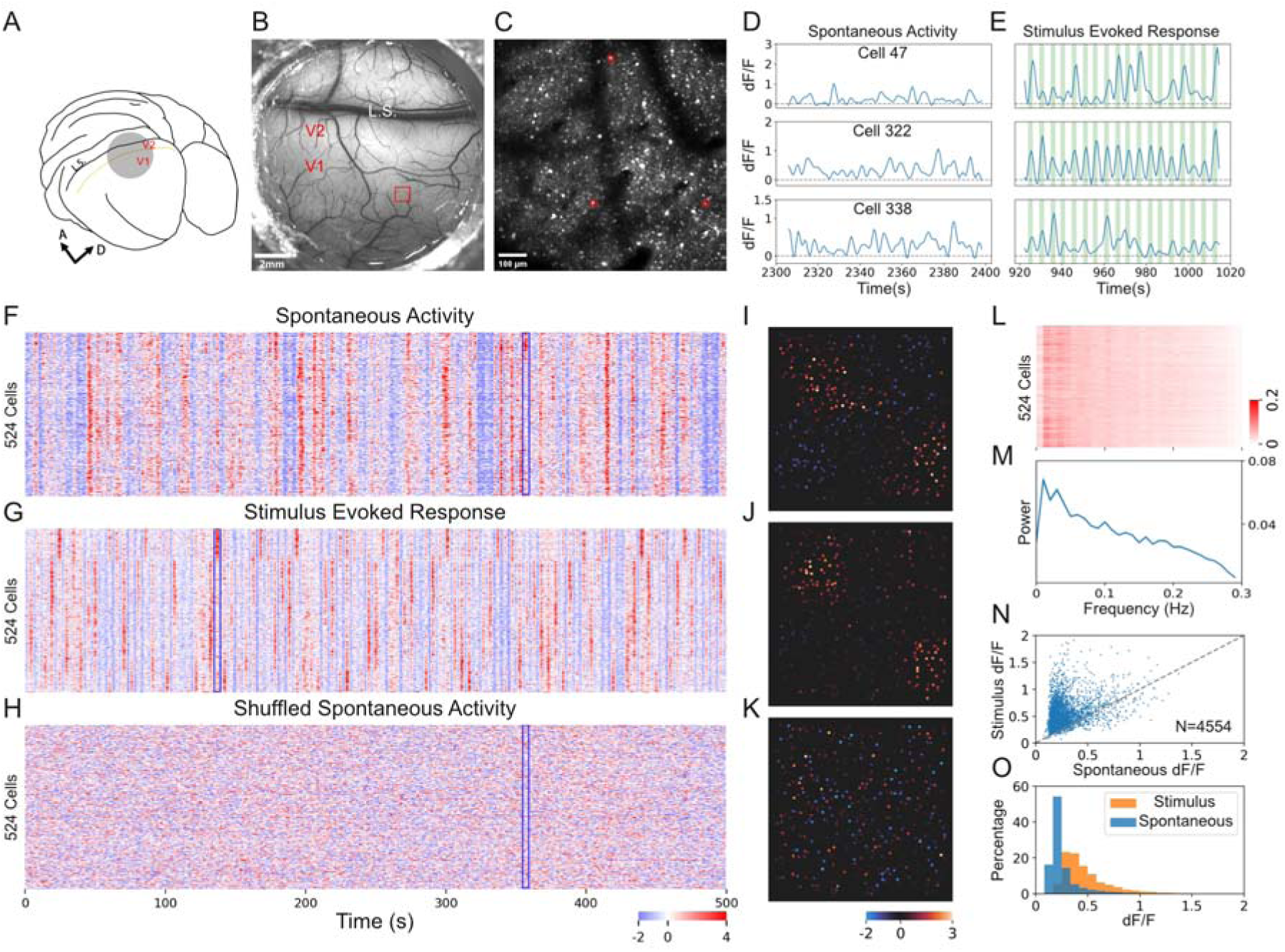
Two-photon calcium imaging of spontaneous activity. **A**. An illustration of the macaque brain and the imaging region (circle) over areas V1 and V2. A: anterior, D: dorsal. L.S.: Lunate sulcus. **B**. In vivo image of a 13-mm diameter optical chamber. The red frame indicates the size of a two-photon imaging site in V1. **C**. Fluorescent image of neurons in the red-frame region in panel B. This 830X830 μm frame was imaged using a 16X objective at a 197 μm depth from the cortical surface. Three neurons marked in red are examined in detail in panel D & E. **D**. Representative spontaneous activity (dF/F) of the 3 example neurons marked in C. Individual neurons showed random activity. **E**. Representative responses to visual stimuli of the same example neurons shown in D. Vertical green bars indicate the stimulus presentation time. The visual stimuli were gratings of various orientations. **F**. Spontaneous activity of all neurons (N=524) imaged in site 1 over a period of 500 seconds. Neurons’ activity was individually normalized. Example frames (blue outline) were examined in detain in panel I. **G**. Responses of the same group of neurons in F to gratings presented at various orientations. **H**. The same spontaneous data as in F but were phase-shuffled for each individual neuron. **I**. An average of 6 consecutive frames (outlined in F) showing the Z values of all neurons at that time point. Only the neurons were shown in this frame. The active neurons exhibited a spatial pattern which was similar to one obtained in stimulus runs. **J**. An average of example frames (outlined in G) extracted from a stimulus run showing the pattern of co-activating neurons. **K**. Example frames (outlined in H) from the shuffled data did not exhibit a meaningful modular pattern. **L**. Frequency spectrums of all neurons in this site. Signals below 0.005Hz and above 0.3Hz were removed in data preprocessing. **M**. Average power spectrum exhibited major signals under 0.1Hz. **N**. Average amplitudes of responses to stimulus and average amplitudes of spontaneous activity for all the neurons imaged in 8 V1 sites (N=4554). **O**. Distributions of response amplitudes for stimulus-driven (orange) and spontaneous activity (blue).

In separate experiment, intrinsic signal optical imaging was also performed for these chambers and basic functional maps were obtained for comparison with the functional maps obtained with two-photon imaging (Figure S1).

### Basic features in spontaneous activity

In single neurons, spontaneous calcium signal appeared to fluctuated randomly (Figure 1D), with an amplitude smaller than their visually driven responses (Figure 1E). Population-wise, neurons showed burst-like co-activation (Figure 1F). These bursts usually had different neuron participants, and some spontaneous frames had similar spatial patterns with the stimulus ones (i.e., functional maps, Figure 1 I&J). This suggests that the spontaneous ensembles were not formed randomly, rather, they were formed by some internal mechanisms that resembled the stimulus effects. As a comparison, after each neuron’s spontaneous activity was phase-shuffled, the overall pattern was completely different (Figure 1H) and no such spatial pattens could be found (e.g., Figure 1K). Single-neuron-wise, their spontaneous activation rates were similar, in a range of 0.01∼0.05Hz (Figure 1 L&M). Their average fluctuation amplitude was about half of that under visual stimulation (Figure 1 N&O). Population-wise, spontaneous ensembles were dynamically formed at a mean frequency of 0.14 Hz, each ensembles lasted ∼4 seconds (Figure S2D & E). Ensemble sizes (number of neurons co-activated) and frequencies (how frequent that size of ensemble observed) were negatively correlated (Figure S2C).

### Orientation patterns in population activity

PCA (Principal Component Analysis) is a useful data-driven analysis tool and frequently used in fMRI data analysis (Carbonell et al. 2011). For the high-order spontaneous data, we used PCA to reduce its dimensionality. As illustrated in Figure 2A, PCA reduced the dimension in cell axis. After PCA, the top 10 principal components (PCs) explained 49.6±6.1% of the total variance (averaged from all 8 V1 sites, Figure 2C), indicating the effectiveness of this procedure. Figure 2B shows the PC space and spontaneous data from an example site, which contained 5352 frames collected in ∼70 min. Each data point represents a frame. The data points were mainly clustered into one cloud, without separate clusters. This suggests that there were no separate dominate states in the imaging period. After dimension reduction along the cell axis, each PC can be viewed as a hypothetical cell, whose activity was a weighted sum of the activity of the original neurons. Each PC thus can be visualized as a frame of the cell weights (e.g., Figure 2D). PC1 appeared to be a global pattern, in which all neurons had a high weight. Indeed, its timecourse was almost identical to the sum of all neurons’ z-scored activity (Figure S3B). It accounted for 35.1±5.1% variance in the data, indicating that the co-activation of neurons in the whole imaging region was the most dominant component in the spontaneous activity. The rest of PCs explained much less of the total variance, but each had a spatial pattern, some of them resemble the orientation map in that region (Figure 2 E&F), others had a better correlation with ocular dominance or color maps (Figure 2E). These correlations were significantly higher than those obtained with shuffled data (Figure 2G).

**Figure 2.**
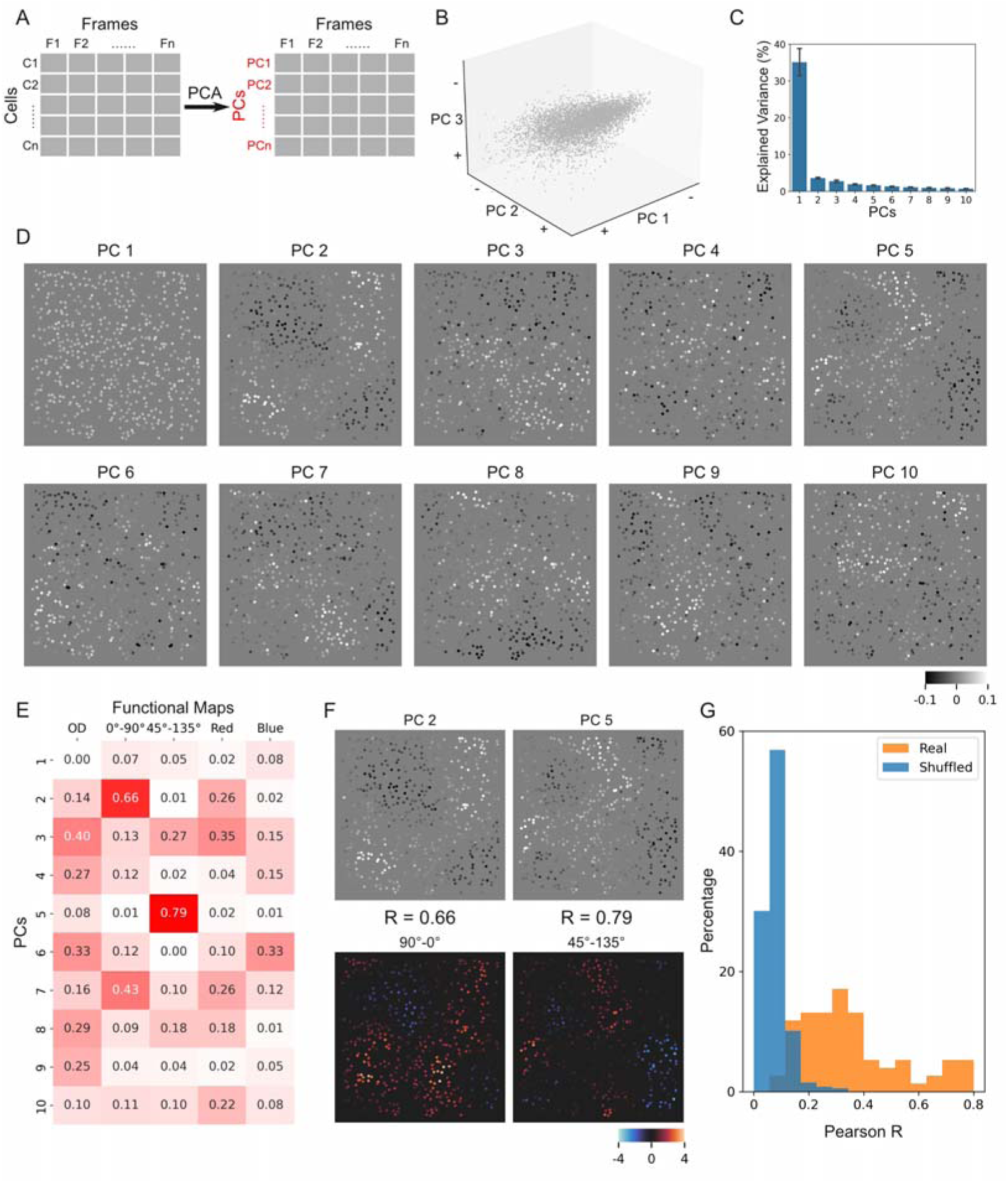
Spontaneous PCA results in cell space. **A**. Illustration of spontaneous data structure and PCA. Spontaneous activity data were organized into two-dimensional matrix: cell dimension and frame dimension. PCA was then carried out in the cell dimension and resulted hypothetical neurons as PCs (cell PCs). **B**. All spontaneous frames from one example site plotted in the PC space. **C**. Percentages of the response variance accounted for by the top 10 PCs. **D**. Visualization of the top 10 PCs, each was a weight map representing the different weights of the original neurons on this PC. **E**. Correlation matrix between the PCs and functional maps of the site. Many PC had similar patterns with the functional maps. **F**. Two PCs (same as shown in D) having the highest correlation scores (in E) are shown (top) along with their best-matching functional maps (bottom). **G**. Distribution of all the correlations between PCs and functional maps (blue, calculated from all 8 V1 sites), compared with the distribution of the same correlations calculated for shuffled data (orange). Each PC contributed one correlation value (the best correlation), thus total 80 correlations were plotted (real). For the shuffled data, each spontaneous data was shuffled 500 times, and 80 correlation values were obtained for each shuffle.

As a control, we phase shuffled each neuron’s spontaneous activity (the same as in Figure 1H), and did the same PCA on the shuffled data. The result PCs had a very low power in accounting for the total variance (top 10 PCs explains a total of 4.1±0.7% of the variance, Figure S4B). Neither did the visualized PCs have systematic pattern or correlate with functional maps (Figure S4 C&D). This supports that the patterns we observed in spontaneous data were non-random in nature.

Since the PC axes contained orientation features (Figure 2 E&F), we can verify this by plotting orientation response data into this PC space. We plotted cells’ responses to gratings (new data) into the PC space obtained in spontaneous data analysis (old space). As we can see, frames obtained with 8 different grating orientations (different colors) clustered into 8 different branches with neighboring branches representing neighboring orientations (Figure 3B). This proves that the space is indeed organized with respect to the orientation information.

**Figure 3.**
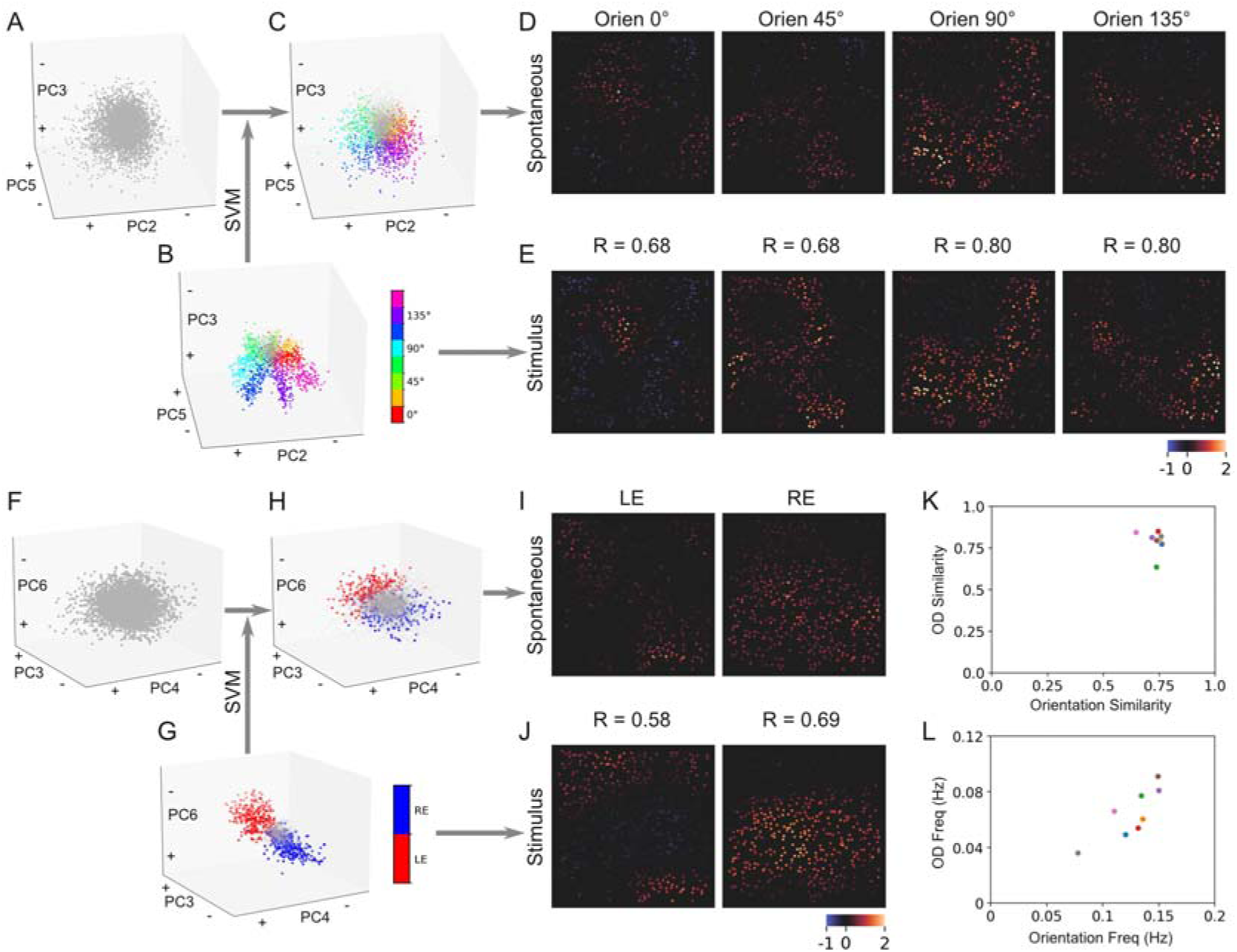
Spontaneous orientation and OD patterns in V1. **A**. Spontaneous frames in the PC space shown with axes of PC2, PC3, and PC5. This case is the same as the one shown in Figure 2B. **B**. Stimulus-driven frames plotted in the PC space resulted from PCA on spontaneous data (i.e., the same space as in A). Gray dots are frames imaged during inter-stimulus-intervals (ISI). The coordinates of the frames were used to train a SVM classifier to categorize stimulus orientations (different colors). **C**. The same plot as in A except that frames were colored based on the SVM classification of these spontaneous frames. Different colors represent different orientation categories that are the same as in B, and gray ones were classified as non-orientation category (ISI frames). **D**. Averages of the same-category frames in C. These maps had high correlations with those functional maps obtained from stimulus-driven images (E). **E**. Stimulus-driven orientation maps (single-condition maps). **F-J**. The same analysis as in A-E, but on ocular dominance features. **F**. The same spontaneous frames in the PC space shown with axes of PC3, PC4, and PC6 (same data as in A). **G**. Stimulus-driven frames collected in monocular stimulation runs plotted in the spontaneous PC space (same space as in F). Red, Blue and gray dots represent left-eye, right-eye, and ISI frames, respectively. Their coordinates were used to train a SVM classifier to separate left-eye, right-eye, and ISI frames. **H**. Spontaneous frames as in F and colored according to the SVM classified categories. **I**. Left-eye (LE) and right-eye (RE)-patterns obtained from the classified spontaneous frames. They had a good correlation with the functional maps in J. **J**. Left- and right-eye functional maps obtained from stimulus-driven frames. **K**. Orientation and ocular dominance correlation values, as shown in D&E and I&J, for 8 imaged V1 sites. These values were similar across different V1 sites. **L**. Occurrence frequencies of orientation ensembles and ocular dominance ensembles, which were correlated across 8 imaged V1 sites.

Now with the coordinates of orientation frames in the PC space, we can further examine whether spontaneous frames can be classified into different orientation groups. We trained a SVM classifier using the PC coordinates of the orientation frames (Figure 3B). We only used the coordinates of the first 10 PC axes and achieved 86.0% correct rate (5- fold cross validation) in the 9-category classification (8 orientations and 1 blank). We then used this classifier to examine the spontaneous frames. There were 25.1% frames that were classified as orientation frames, the rest (74.9%) were classified as blanks (Figure 3C). Compared with the separate branches in stimulus frames (Figure 3B), the spontaneous frames were more continuous (Figure 3C), indicating a more uniform orientation representation in the spontaneous activity. PC 2, 3, and 5 had the largest contribution in the SVM classification, therefore were used to visualize the PCA results (Figure 3 A-C). Finally, the averaged spontaneous frames in each orientation category were very similar to those obtained from stimulus frames (Figure 3 D&E), and had higher correlation values than the controls obtained with artificial stimulus data (Figure S5). The correlation was around 0.7, which further confirms the existence of orientation-specific co-activation in these neurons. Although PC1 represented global activity and did not contribute to specific orientation patterns, frames had higher PC1 weights tended to have higher chance to be classified as an orientation pattern (Figure S3 D&E).

### Ocular dominance and color patterns in spontaneous activity

During each experiment, we also collected neurons’ responses to monocular black/white grating, as well as responses to binocular color gratings. We used the same approach and plotted these stimulus-driven data (frames) into the old PC space obtained with spontaneous data (i.e., same as Figure 2B). Figure 3G shows that the frames collected during left-eye stimulation (red) and right-eye stimulation (blue) were well separated in the PC space. A SVM classifier trained with the coordinates of these frames subsequently identified 10.0% of the spontaneous frames as monocular activation frames (Figure 3H). The results were confirmed by the similarity between the averaged spontaneous frames and left- or right-eye functional maps (Figure 3 I&J).

We also examined whether color response patterns existed in the spontaneous activation. With the same method, we found some spontaneous frames can also be classified as similar to those obtained in color grating stimulation (Figure S6C). However, the averaged color spontaneous patterns had weaker similarity with the color functional maps (Figure S6 D&E), the correlation values ranged from 0.18 to 0.63(Pearson’s r =0.43±0.1 overall). We conclude that although co-activation of color-selective neurons did exist in spontaneous activity, it was not as robust as those for orientation and ocular dominance ones. This is consistent with our previous findings obtained with intrinsic optical imaging (Cai et al. 2023).

So far, spontaneous patterns were only exhibited in form of neurons (Figure 3 D&I, Figure S6D). We also examined corresponding spontaneous patterns in form of frames (Figure S7), in which original dF/F frames of the classified frames were averaged. Although the correlation scores were lower, the overall similarity was evident for corresponding spontaneous patterns and functional maps (Figure S7).

From each of 8 V1 imaging sites, we obtained clear orientation and ocular dominance patterns in spontaneous activity. These patterns had an average correlation value of 0.8 with corresponding functional maps. Variation among imaging sites were small (Figure 3K). As a comparison, correlation value for color pattern is 0.43. The occurrence frequency of these patterns varied from experiment to experiment (Figure 3L), which may due to different anesthetic statues of the animals in each experiment. The frequencies for orientation and ocular dominance in one experiment were normally correlated (Figure 3L), which is consistent with this hypothesis. In summary, these observations indicate that neurons with common preferences co-activated not only during visual stimulation, but also in ongoing activity.

### Correlated activity among similarly tuned neurons

So far, we did PCA dimensional reduction in *cell* dimension. Similarly, we did PCA in *frame* dimension in order to see whether cells may cluster based on this data-driven approach. As shown in Figure 4A, each PC axis represents one frame, and each data point in the frame space is a cell. The dimensionality of this frame space was higher, as the top 20 PCs only explained 31.9±4.2% variance (Figure 4B). However, we observed a separation of different orientation-preferring neurons (Figure 4C), as well as separations of left-eye and right-eye preferring neurons in this PC space (Figure 4D). This indicates that neurons with common tuning preferences had similar spontaneous activation, and that these tuning features (ocular dominance, orientation) were among the top factors that influenced neurons’ spontaneous activity (among the top PCs).

We calculated average vectors for neurons preferring 0° and 90° and vector for neurons preferring 45° and 135° (arrows in Figure 4C). Visualization of these vectors exhibited orientation patterns (Figure S8A&B). These two vectors were orthogonal to each other in this high-dimensional PC space (Figure S8D). In addition, these two vectors were both orthogonal to the vector for ocular-dominance neurons (arrow in Figure 4D, also see Figure S8D). These observations indicate that these factors had orthogonal influences on neurons’ spontaneous activity, and different functional networks had independent activity, did not affect each other.

**Figure 4.**
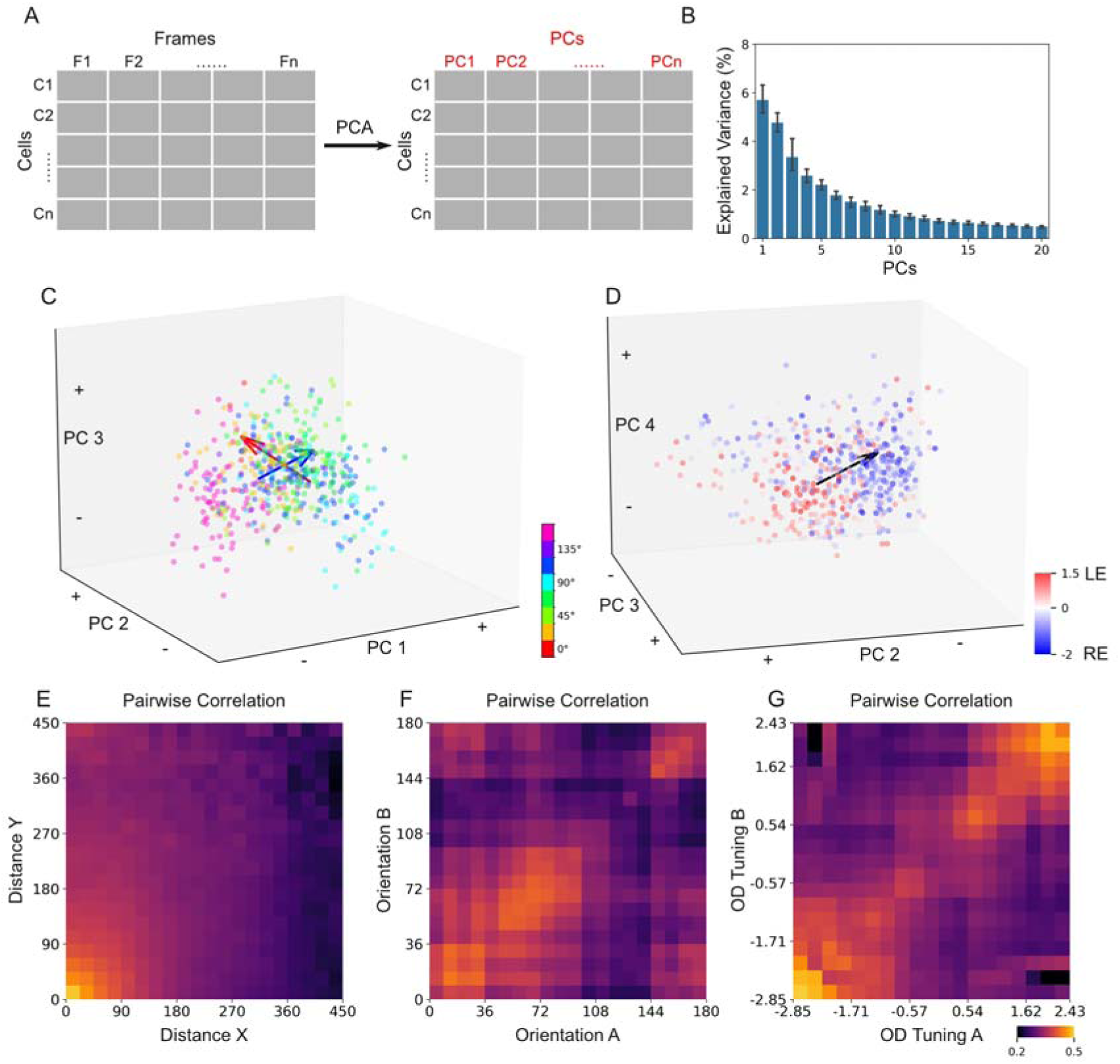
Spontaneous PCA results in frame space. **A**. Similar to Figure 2A, but PCA was carried out along the frame dimension and resulted hypothetical frames as PCs (frame PCs). **B**. Percentages of the response variance accounted for by the top 20 PCs. **C**. Neurons in the PC space colored according to their orientation preferences. Red arrow is a 0° vs. 90° vector, which starts from the mean location of all the 90°-preferring neurons and points toward the mean location of all the 0°-preferring neurons. Similarly, blue arrow represents the 45° vs. 135° vector. These two vectors are orthogonal to each other. **D**. Same plot as in C but colored according to neurons’ ocular dominance preferences. The black arrow represents the LE vs. RE vector, which is also orthogonal to the orientation vectors shown in panel C. **E**. Pairwise correlations between the responses timecourses of any two different neurons. The correlation values were sorted and averaged according to the distances of the two neurons. **F**. Same as in E, but sorted with neurons’ orientation preferences. **G**. Same as in F and E, but sorted with neurons’ ocular dominance preferences.

These results can also be directly obtained by calculating pair-wise correlations between neurons’ spontaneous activity. Although most neurons were positively correlated (due to the global signal), the degree of correlation varied. Figure 4E shows that correlation dropped when the distance between neurons increased. This might due to the functional clustering. Figure 4F shows that neurons with similar orientation-preferences had stronger correlation. Figure 4G shows that neurons with similar eye-preferences had stronger correlation. Neurons with weak eye-preferences (binocular neurons) or neuron pairs had different eye preferences had weaker correlations.

### Spontaneous activity in V2

We also imaged 3 sites in area V2. The 3 sites sited in different locations with respect to V2 stripes, as indicated by functional maps obtained with intrinsic signal optical imaging (Figure S1). Unlike V1 neurons, neurons in V2 are mostly binocular and have no/weak eye preferences. In addition, different stripes exhibit large differences in orientation selectivity and color selectivity. We did the same analysis for the V2 data. Figure 5A shows the data from a site located in a thick/pale stripe. This site had strong orientation selectivity and weak color selectivity. Accordingly, orientation patters obtained from its spontaneous activity was strong (Figure 5E), but color was weak (not shown). Figure 5F shows data from another V2 site, which was located in a thin stripe. The spontaneous activity in this site exhibited strong color patterns (Figure 5J). The correlation value (∼0.6) was higher than all those observed in V1 (Figure 5K).

**Figure 5.**
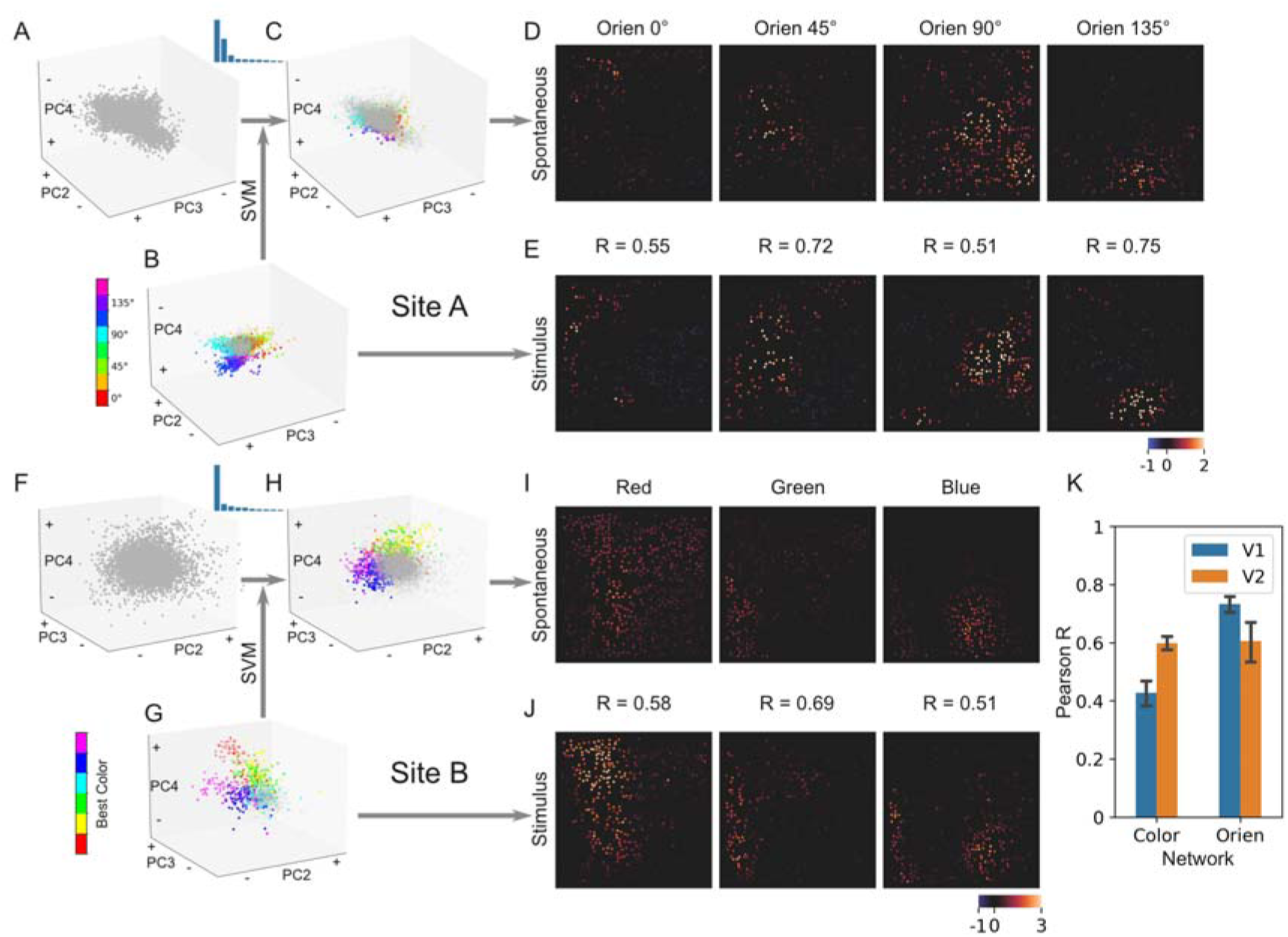
Spontaneous orientation and color patterns in V2. **A-E**. Similar to the PCA and SVM classification procedures shown in Figure 3, but for a V2 site. Similar to V1, spontaneous orientation patterns were observed (E). Bar plot represent top 10 PC’s explained variance ratio. **F-J**. Similar to above, but for another V2 site. This V2 site had weak orientation selectivity, but strong color selectivity. The spontaneous color patterns (K) were observed and well correlated with functional maps (L). **K**. Correlation values between spontaneous patterns and functional maps for all 3 V2 imaging sites (orange) compared with those from V1 (blue).

Thus, like in V1, spontaneous activity in V2 also exhibited patterns that were correlated with the underlying functional architectures. There were also differences between these two areas. For example, V2 had stronger color spontaneous patterns than V1. In addition, 3 imaged V2 regions had very different features, due to the small imaged regions and large V2 functional architectures. In V1, the 8 imaged regions had similar properties, indicating that the imaged regions had a scale close to the functional element of V1, the hypercolumn.

## Discussion

We analyzed spontaneous activity of population neurons in V1 and V2 in anesthetized macaques. We found that neurons dynamically formed different active ensembles. The ensembles resembled the known functional architectures, including different orientation-preference patterns, ocular dominance and color patterns. As a result, neurons had common selectivity tended to have higher correlated activity. In V1, orientation and ocular dominance both had strong effect on neurons’ spontaneous activity, while color’s effect was weaker. In V2, depending on the particular locations, orientation and color could had strong or weak effects on neurons’ spontaneous activity. These results provided direct evidence for the hypothesis that spontaneous activity depends on underlying functional architectures. Different architectures have different contributions. Further, the neuronal evidence supports previous findings obtained with voltage-sensitive-dye imaging (Omer et al. 2019) and hemodynamic imaging (Cai et al. 2023) and proved the neural origin of the mesoscopic functional connectivity they observed.

### Preferential horizontal connection model for patterned spontaneous activity

What we observed can be explained by non-uniform horizontal connections. Previous studies have demonstrated preferential horizontal connections among neurons having similar tuning properties, including orientation preferences (Gilbert & Wiesel 1989; Kisvarday et al. 1997), ocular dominance (Malach et al. 1993; Yoshioka et al. 1996), and color preferences (Livingstone & Hubel 1984; Yoshioka et al. 1996; Yabuta & Callaway 1998). These domain-specific connections coexist with a large amount of domain-non-specific connections, which connect opposite selective neurons (Malach et al. 1993; Yoshioka et al. 1996). It has been proposed that it is “like-to-like” connections within short cortical distances (e.g., within a hypercolumn), and “like-to-all” connections for longer distances (Chavane et al. 2022). These horizontal connections are the morphological basis for the correlated ongoing activity.

As illustrated in the conceptual model in Figure 6, the domain-non-specific connections (gray lines in Figure 6B) are the morphological basis for the global activity we observed (PC1 in Figure 2); and domain-specific connections among different types of functional domains (blue and red lines in Figure 6B) are the basis for the multiple spontaneous patterns (e.g., PC2 in Figure 2). A single V1 neuron normally located in the junction of multiple types of networks (OD, orientation, color) and fires when it receives enough inputs from earlier level and horizontal connections. Its activation also has non-uniform impact with its surrounding neurons through these non-uniform connections. Two types of connections are normally co-activated, as we have shown that the strengths of PC1 and PC2 are correlated (Figure S3D). It has been shown that, comparing to spontaneous fluctuations around a single background state, such multi-state spontaneous activity is often slow and low-dimensional (Goldberg et al. 2004), which is consistent with our observations. Besides horizontal connections, spontaneous activity in LGN will also affect V1 activity, as well as those from top-down feedbacks. For simplicity reasons, these factors are not included in the model shown in Figure 6.

**Figure 6.**
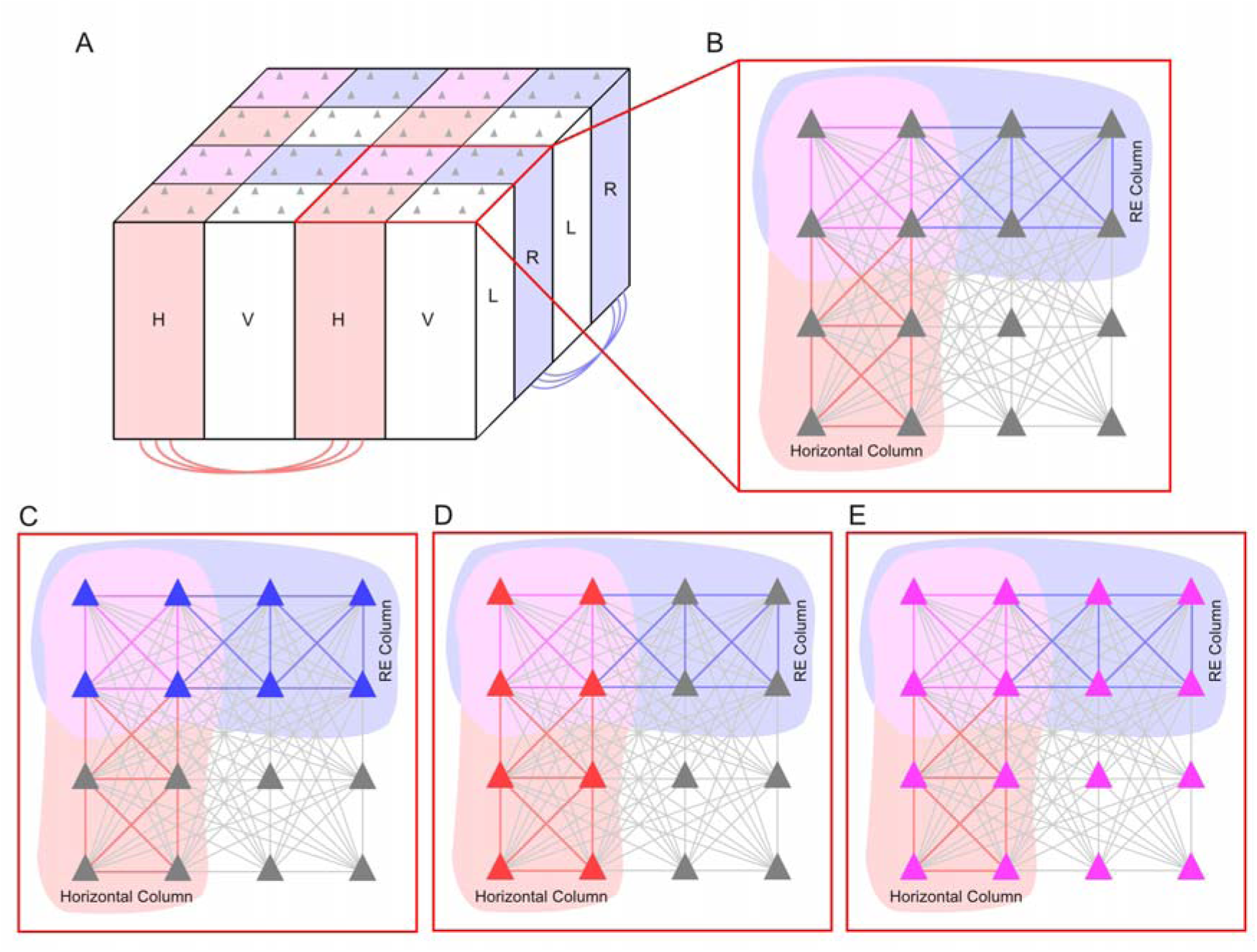
Horizontal connection model. A. A simplified hypercolumn containing left- and right-eye columns (L, R), and horizontal and vertical orientation columns (H, V). Triangles represent neurons. Red and blue lines illustrate domain-specific connections that preferentially connect domains with common selectivity features. B. Surface view of a hypercolumn. Different types of horizontal connections are illustrated, including domain-specific connections among right-eye-preferring neurons (blue lines), horizontal-orientation-preferring neurons (red lines), and domain-non-specific connections (gray lines). C. Coactivation of right-eye-preferring neurons (blue triangles). D. Coactivation of horizontal-orientation-preferring neurons (red triangles) E. Coactivation of all neurons within the hypercolumn (pink triangles).

### Spontaneous patterns as a common phenomenon

Two-photon calcium imaging has been used in studying spontaneous activity in many species. For example, in mice (Goltstein et al. 2015; Mizuno et al. 2018) and rat (Ch’ng et al. 2010), it has been shown that neurons with common orientation preference tend to synchronize. In ferrets (Smith et al. 2018; Mulholland et al. 2021) and cats (Ch’ng et al. 2010), spontaneous orientation maps have been observed. In this study, we observed spontaneous ocular-dominance and color maps, in addition to orientation maps. Thus, this appears to be a cross-species common feature.

Correlations between spontaneous activity and functional architectures were also observed in other sensory cortices. For example, multi-electrode recordings have shown such evidence in rat auditory cortex (See et al. 2019); ECoG recordings in macaque area STP revealed a tonotopic organization in spontaneous activity (Fukushima et al. 2012); intrinsic signal optical imaging of spontaneous activity in somatosensory and motor cortices also revealed functional connections (Card et al. 2022). In higher level areas, for example PFC, spontaneous activity has revealed sub-regions (Kiani et al. 2015). Thus, the dependence of spontaneous activity on functional architecture has been observed in different areas, and in different species. Since these two factors are highly correlated, we suggest that unknown patterns in spontaneous activity could serve as an indicator for new functional architectures. This has important methodological meaning for studying high level areas, which are usually too complex to be studied with classical stimuli or tasks.

### Methodological Comparisons

The present results support our previous findings obtained with intrinsic signal optical imaging (Cai et al. 2023). Both studies used the similar experimental procedures. Both found 3 types of spontaneous patterns in V1, in which orientation and ocular dominance patterns are strong, and color ones are weaker. In the two studies, spontaneous events have similar width (intrinsic signal: ∼4 sec, calcium signal: 4.1 sec in Figure S2D), and temporal frequencies (intrinsic signal: 0.02-0.03Hz, calcium signal: <0.1Hz in Figure 1 L&M). Thus, the present results support the neural origin of FC observed with hemodynamic signal, as well as the usefulness of such signals in study mesoscopic FC (Vasireddi et al. 2016; Card et al. 2022, Cai et al. 2023).

Functional MRI revealed large-scale, long-distance FC in the brain (Fox & Raichle 2007). Limited by its spatial resolution, fMRI is unable to detect sub-millimeter FCs, such as FCs between V1 orientation modules. Optical imaging methods such as VSD, ISOI and wide-field fluorescent imaging are useful in revealing FCs in the mesoscopic scale, but are unable to correlate with single-neuron activity. Two-photon imaging provides a good combination of both spatial and temporal resolutions, makes it an important tool in study fine-scale FC. Our results not only showed mesoscale FCs, but also revealed a global signal (PC1) that all neurons in the field of view contributed. This global signal had a slow temporal frequency (<0.1Hz, Figure 1 L&M), which may correspond to the infra-slow activity observed in fMRI (Fox & Raichle 2007) or wide-field calcium imaging (Mitra et al. 2018), providing a cellular-level support for the resting state BOLD signal.

### Limitation of the present study

This study was carried out on anesthetized animals. Although it has been shown that there are similar FC patterns in both anesthetized and awake animals (Vincent et al. 2007), some studies have also revealed that in anesthetized animals, spontaneous patterns were stronger (Omer et al. 2019). In addition, the type of anesthesia drug may also affect the strength of FC observed (Mura et al. 2022). Recent large-scale recordings in awake animal revealed more complex (higher dimensional) spontaneous activity, some may be related to animal behavior (review in Avitan & Stringer 2022). These behavioral or cognitive factors are more variable and add additional complexity into spontaneous signals. Spontaneous signals in anesthetized animals are simpler and easier to analyze and provided a baseline for further investigations. Signal-wise, the calcium signals we examined were slow and contained sub-threshold neuronal activity, which were different from spike signals. Finally, the PCA analysis detects the common activity in neuron population, thus single neuron characteristics were not examined in this study.

## Materials and Methods

### Animals

Three adult macaque monkeys (two *Macaca mulatta*, one *Macaca fascicularis*, Beijing Inst. of Xieerxin Biology Resource and Hubei Topgene Biotechnology Co., Ltd.) were imaged in this study. All procedures were performed in accordance with the National Institutes of Health Guidelines and were approved by the Institutional Animal Care and Use Committee of the Beijing Normal University (protocol number: IACUC(BNU)- NKCNL2016-06).

### Surgery procedures

Animals were sedated with ketamine (10 mg/kg) or Zoletil 50 (tiletamine HCl and zolazepam HCl, 4 mg/kg). They were placed on a stereotaxic and artificially ventilated with isoflurane (1.5–2.5%). A 22–24 mm diameter circular craniotomy and durotomy were performed in order to expose portions of areas V1 and V2. The craniotomy center was 18 mm from the midline, 14 mm from the posterior bone ridge (Figure 1A).

In each case, we injected AAV9.Syn.GCaMP6S.WPRE.SV40 (#100843, titer ≥ 1×10¹³ vg/mL, Addgene) into 10–15 cortical locations (520 nL each location) at a depth of 600- 800 μm. Cortex was then protected with a piece of artificial dura, the retracted dura was covered back and glued to the top of the artificial dura with medical adhesive. The original bone piece was placed back, secured with a titanium mesh and bone screws. The scalp was sutured. After at least 8 weeks of virus expression, a second surgery was performed. The old craniotomy was reopened, a second round virus injections were performed, followed by implanting of an optical titanium chamber (inside/outside diameters: 13/15 mm).

### Two-photon Calcium Imaging

After the animal recovered from surgery, two-photon calcium imaging was performed every 7-10 days. Animal preparation was the same as those in the surgery experiments, except that animal’s head was tilted for ∼45° to align with the fixed vertical imaging axis. Anesthesia was switched from isoflurane to i.v. infusion of a mixture of propofol (2 mg/kg for induction and 2 mg/kg/hr for maintenance) and sufentanil (0.15 μg/kg for induction and 0.15 μg/kg/hr for maintenance). The animal was immobilized with vecuronium bromide (0.25 mg/kg for induction and 0.05-0.06 mg/kg/hr for maintenance) to prevent eye movements. Anesthetic depth was assessed continuously via monitoring the electrocardiogram, end-tidal CO_2_, blood oximetry, and body temperature. In order to have a stable anesthesia level, imaging usually started more than one hour after the anesthesia switch.

In each imaging experiment, spontaneous imaging session was always performed before the visual stimulus sessions. The monkey’s eyes were closed and covered with an additional eye-cover. The excitation wavelength of the laser (generated by Chameleon Ultra II, Coherent Inc) was set at 980 nm. Under a 16X objective (0.8 N.A., Nikon), 515X512 pixel images, covering an 830X830 μm cortical surface, were collected continuously at a frame rate of 1.3 Hz (Brucker Ultima IV Extended Reach, Bruker Nano Inc). Image plane was at a depth of 180-270 μm. Spontaneous imaging session usually lasted for 1-2.5 hr, during which we tried to maintain a stable environment with minimum light and sound.

After spontaneous imaging, the monkey’s eyes were opened. Pupils were dilated (tropicamide) and fit with contact lenses of appropriate curvatures to focus on a stimulus screen 57 cm from the eyes. Optical disks were plotted on the screen in order to estimate the approximate locations of the fovea. In visual stimulation runs, images were collected continuously as in the spontaneous runs. Each visual stimulus was presented for 1.5 second and separated by a 0.5 second inter-stimulus-interval. The beginning of each stimulus presentation was synchronized with the beginning of a frame scanning.

During the imaging session, slow drifts of cortex in the imaging field of view was observed. The drift was normally less than 100 μm in the X-Y plane and less than 80 μm along the Z-axis in a course of 6–8 hr. We frequently checked the cell features during the imaging and adjusted position of the focal plane accordingly. Drifts in the X-Y plane were further corrected in offline data analysis.

### Visual Stimuli

Visual stimuli were generated using ViSaGe (Cambridge Research Systems Ltd.) and presented on a 21-inch LCD monitor (Dell E1913Sf) positioned 57 cm from the eyes. The stimulus screen was gamma corrected and worked at 60 Hz refreshing rate. Left- and right-eye visual fields were diverged by a pair of 9° prisms to prevent their overlap on the screen. A black board were placed between the two eyes to prevent cross eye stimulation. For black/white gratings, the luminance for white bars was 100 cd/m^2^. Within each run, the stimuli were presented in a random order.

#### Stimuli for RF mapping

Circular grating patches sized 2.5° were presented at a grid of 5X5 locations. Each grating had a rectangle waveform (duty cycle: 0.2; SF: 1.5 c/deg; TF: 8c/sec) and was presented at 4 different drifting directions (45°, 135°, 225°, 315°). Each stimulus was repeated for 3 times (total 5×5×4×3=300 trials). RF positions were online analyzed and used for following stimulus presentation. Two eyes were mapped separately. The mapped population RF had an eccentricity of 3-5°.

#### Stimuli for estimating neuron’s eye dominance

A 7°X7° grating patch was presented monocularly to the left or right eye. The mapped population RF locations for the two eyes were used here for the center locations of the square grating patch. Gratings are rectangle wave (duty cycle: 0.2; SF: 1.5 c/deg; TF: 8c/sec). Stimuli were presented in a block design. Each block contains 4 orientations (0°, 45°, 90°, 135°), two locations (for two eyes), as well as a blank condition (total 9 conditions). Gratings were drifting at random directions orthogonal to the gratings. Normally 20-30 blocks were presented.

#### Stimuli for estimating neuron’s orientation preference

Similar to eye-dominance stimuli, grating patches were used for orientation test, except that the left- and right-eye stimuli were presented together through a full screen patch. Each block contains 17 conditions, including 8 orientations (in 22.5° steps), each drifting at two opposite directions, and a blank condition.

#### Stimuli for estimating neuron’s color preference

Color stimuli were sinewave gratings (SF = 0.15c/degrees, TF = 1c/s) in full screen. Each block had 29 conditions, including 1 blank condition, and gratings in 7 different colors, each presented at 4 orientations (0°, 45°, 90°, 135°), drifting at one of the two directions randomly chosen orthogonal to the gratings. The 7 colors were: red (255,0,0), yellow (255,255,0), green (0,255,0), cyan (0,255,255), blue (0,0,255), purple (255,0,255) and white (255,255,255). The mean luminance for white gratings was 50 cd/m^2^, blue gratings were 7 cd/m^2^, and all other color gratings were 20 cd/m^2^.

## Data Analysis

### Cell identification

We first did motion correction and cell identification with CalmAn toolbox v1.9.15 (Giovannucci et al. 2017, 2019) and Python v3.10.8, and obtained timecourses of fluorescent strength for individual neurons.

### Preprocess

Each timecourse were first filtered with a Butterworth filter (5^th^ order, bandpass 0.005- 0.3Hz) to remove slow drifting background and breath noise. Then fluorescent change was calculated as dF/F=(F-F0)/F0, in which F0 is the mean value of the lowest 10% F values in the current run.

We did Z-score normalization for dF/F series before PCA, using 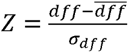, in which *dff* is the dF/F of the current frame, 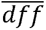 is the mean dF/F of the series, σ*_dff_* is the standard deviation of the dF/F of the series.

### Visual feature preference

Neuron’s ocular dominance was evaluated using Cohen’s D : *Cohen* 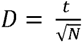, in which t is the t-test value for responses to the 4 left-eye stimuli and 4 right-eye stimuli, N is the number of samples.

Neuron’s responses to 16 oriented gratings were fitted with a modified von Mises function (Mardia 1972): 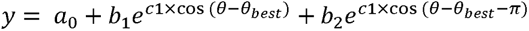. Neurons had R^2^>0.7 were considered had orientation preference and its preferred orientation was then obtained from this fitting function.

Neuron’s color selectivity was evaluated by comparing its responses to each color grating and black-white grating (t-test). A t-value larger than 0 and p<0.05 were considered had color selectivity to that color. Thus, a neuron can have more than one preferred color.

### Functional maps

We obtained functional maps for ocular dominance, orientation, and color for each imaging sites. Each had two types: single-condition maps and difference maps. Use ocular dominance maps as example, there were two single-condition maps: left-eye condition map and right-eye condition map, each was obtained by calculating the mean of the Z-score for each cell. The difference map was a Cohen D map calculated using the Z-scores to left and right eye. For neurons’ functional map, each neuron was simplified as a small disk in its original location and had a value calculated above.

### Shuffled data

We used phased-shuffled spontaneous data as controls: each neuron’s Z-score time series were first Fourier transformed and the phases of between 0.005-0.3Hz were randomized. These frequency data were then transformed back to time series. These shuffled time series maintained the same power spectrum as original ones.

### PCA

PCA (Principal Component Analysis) is a useful data-driven analysis tool and frequently used in fMRI research (Carbonell et al. 2011). For the spontaneous data, we used PCA to reduce its dimensionality in two separate ways. The first is illustrated in Figure 2A, in which the dimension in Y-axis (cell dimension) was reduced. The second is illustrated in Figure 4A, in which the dimension in X-axis (frame dimension) was reduced.

### SVM classification based on PCA results

In order to examine the similarity between spontaneous frames and stimulus frames, we combined PCA and SVM classification. We first did PCA on spontaneous data to reduce the cell dimension. Then we plotted the frames obtained in stimulus conditions into the PC space. We used the coordinates in the first 10 dimensions to train a SVM classifier (scikit-learn toolbox, v1.3.0). Finally, spontaneous frames were classified in this trained SVM classifier.

### Spontaneous maps

Based on the procedures above, we obtained spontaneous frames that resembles the stimulus-driven frames. A continuous sequence of classified frames was considered an “event”. A spontaneous map can be calculated similarly as the one for stimulus frames.

## Acknowledgements

This work was supported by the National Science and Technology Innovation 2030 Major program (2022ZD0204600) and the National Natural Science Foundation of China (32271079, 31625012, and 31530029). We thank lab members J. Lu, Y. Xiao, for providing technical assistance.

## Supplemental information

Figures S1–S8

## SUPPLEMENTAL FIGURES

**Figure S1.**
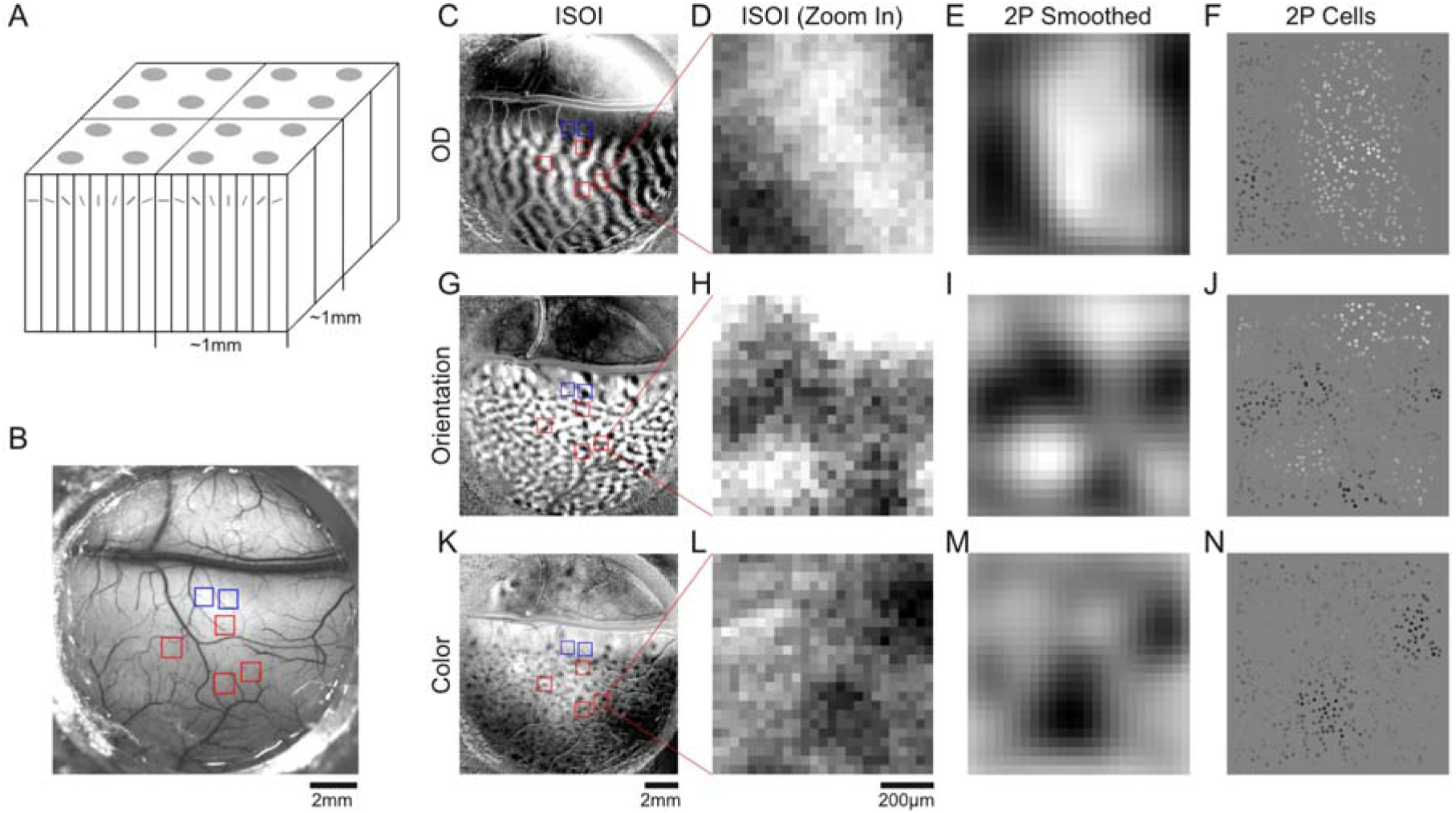
Functional maps obtained with two imaging techniques in Case 1. A. A diagram shows the ice-cube model and hypercolumns. Each hypercolumn covers about 1 × 1 mm cortical surface and contains a set of orientation columns and a set of ocular dominance columns. In this diagram, 4 hypercolumns are shown. B. In vivo image of a 13-mm diameter optical chamber for Case 1. The red and blue frames indicate regions in V1 and V2, respectively, that were subsequently investigated with two-photon imaging. C: Ocular dominance map for the chamber shown in B, obtained with intrinsic signal optical imaging. The two-photon imaging sites (same as in B) were also labeled. D: Zoom-in view of the ocular dominance map (in C) in one of the two-photon imaging site. E: A blurred view of the corresponding functional map obtained with two-photon imaging (shown in F). Maps in D and E had good correspondence. F: Ocular dominance map of neurons in the same region shown in C and D, obtained with two-photon imaging. G-J: Similar to C-F, but for an orientation map of the chamber region (G) and zoomed in region (H-J). K-N: Similar to C-F, but for a color map of the chamber region (K) and zoomed in region (L-N).

**Figure S2.**
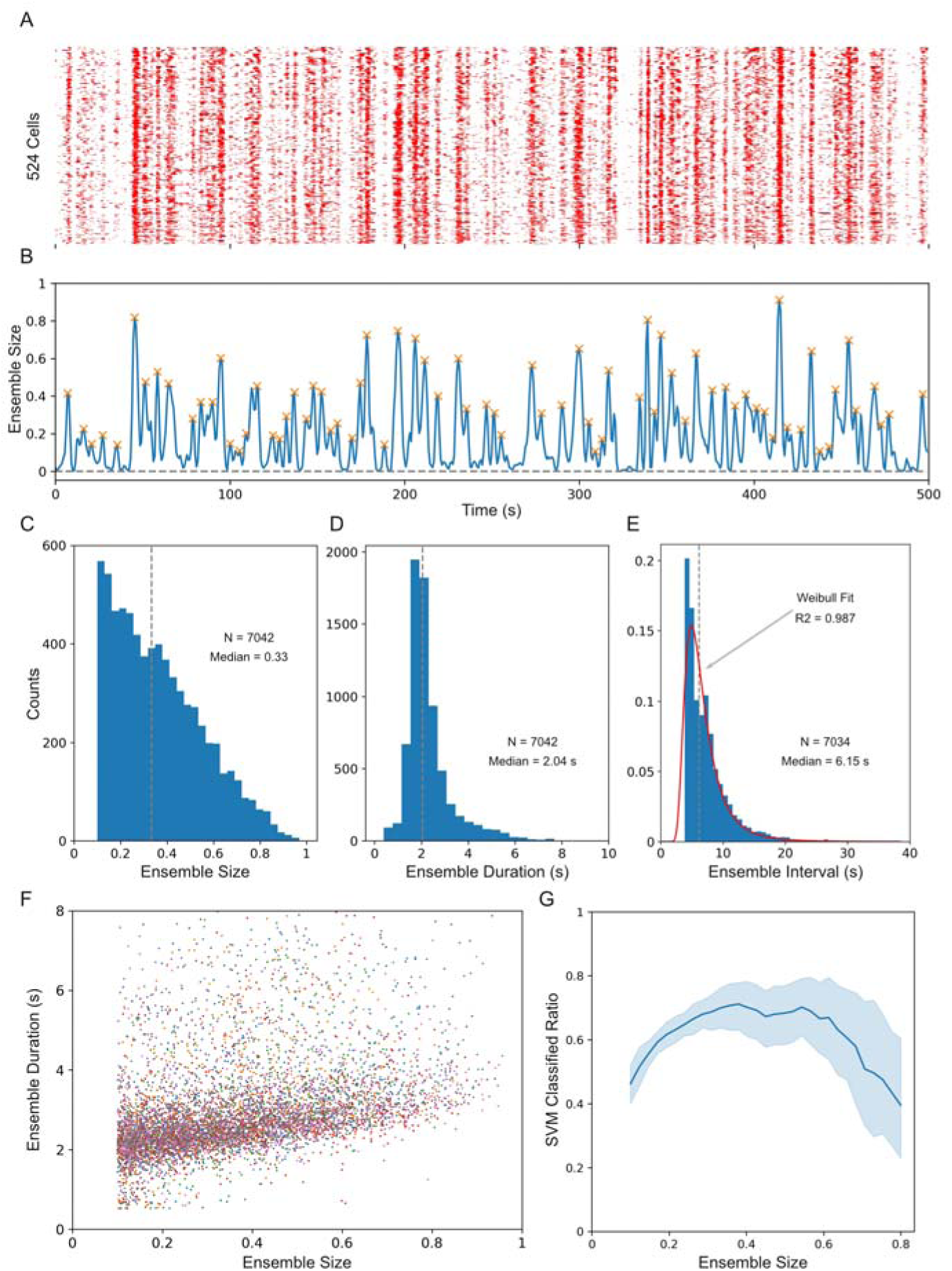
Features of spontaneous ensembles. **A**. Raster plot of spontaneous events for individual neurons over a 500 second period (the same data as shown in Figure 1F). Neuron activity above 1SD of its mean was considered a spontaneous event. **B**. Ratio of activation neurons (as shown in A) in each frame. Frames that had less than 10% of the neurons were not considered, and active frames on both side of a peak were grouped as an activation ensemble (orange crosses). **C**. The distribution of ensemble occurring frequency for different ensemble sizes (how many neurons involved, as a proportion of the total neuron numbers). The overall occurring frequency for all ensembles was 0.140±0.015 Hz. **D**. The distribution of ensemble duration (half height width), the median duration was 4.08 second (half height width: 2.04 seconds). **E**. The distribution of intervals between two ensembles (peaks in B), which can be fitted by a Weibull function (red curve). The vertical dotted line marks the median value of 6.15 seconds. **F**. All ensembles plotted according to their size (X axis) and their duration (half height width, Y axis). Ensembles from different imaging sites were in different colors. It shows larger ensembles were fewer but had longer durations. **G**. Proportions of ensembles that were classified as having a pattern (e.g., colored frames in Figure 3C), plotted in Y axis, as a function of the ensemble size (X axis). It shows that ensembles had medium size (30-60% neurons activated) were more likely to have a spontaneous pattern similar to the functional maps than those smaller and larger ensembles (on the left and right sides of the X axis).

**Figure S3.**
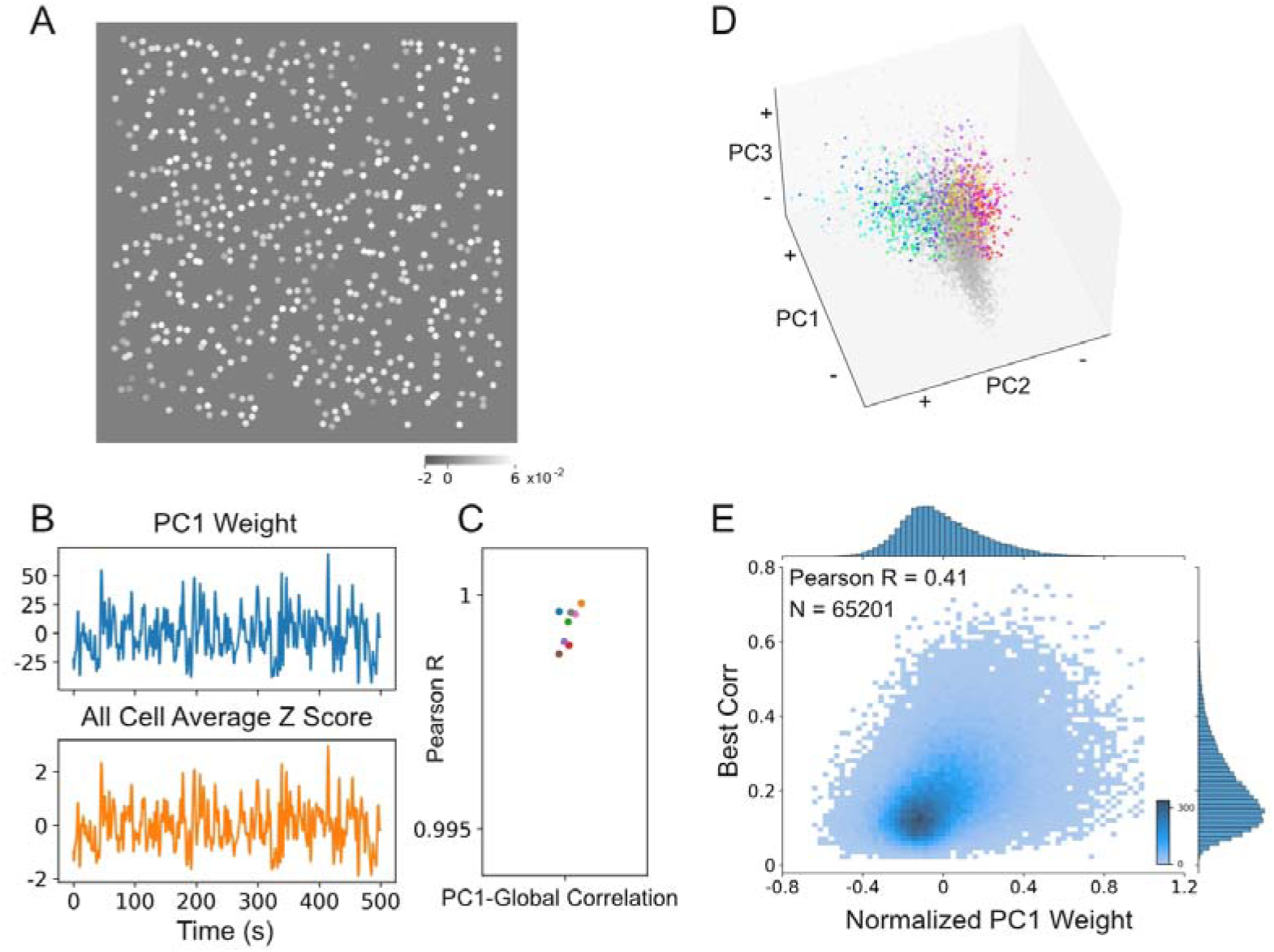
PC1 represent the global activity. A. An example map of PC1 (the same map as shown at the top-left in Figure 2D), obtained with PCA on spontaneous data (the same example case in Figure 2). B. The weight timecourse for the PC1 shown in A (top), and the overall-activity timecourse of the same period obtained by averaging all neurons’ z-scored activity (bottom). These two timecourse had a very high correlation (r>0.99). C. Average correlations for all timecourse pairs as shown in B for 8 V1 sites. D. Frames in the PC space (the same example case as shown in Figure 2B). Colored data points are frames that were identified as orientation frames (as shown in Figure 3C, but with different PC axes). The corn-like shape of the data cloud indicates that when PC1 value increases, PC2 and PC3 values also increase, and more frames were identified as orientation frames. This suggests that when overall spontaneous activity increases, the occurrence of map-like activity also increases. E. The best correlations between each spontaneous frames and functional maps (Y axis) had a positive correlation with the frames’ weights on PC1 axis (X axis). All spontaneous frames (N=65201) from 8 V1 sites were included. This observation is consistent with the trend shown in panel D.

**Figure S4.**
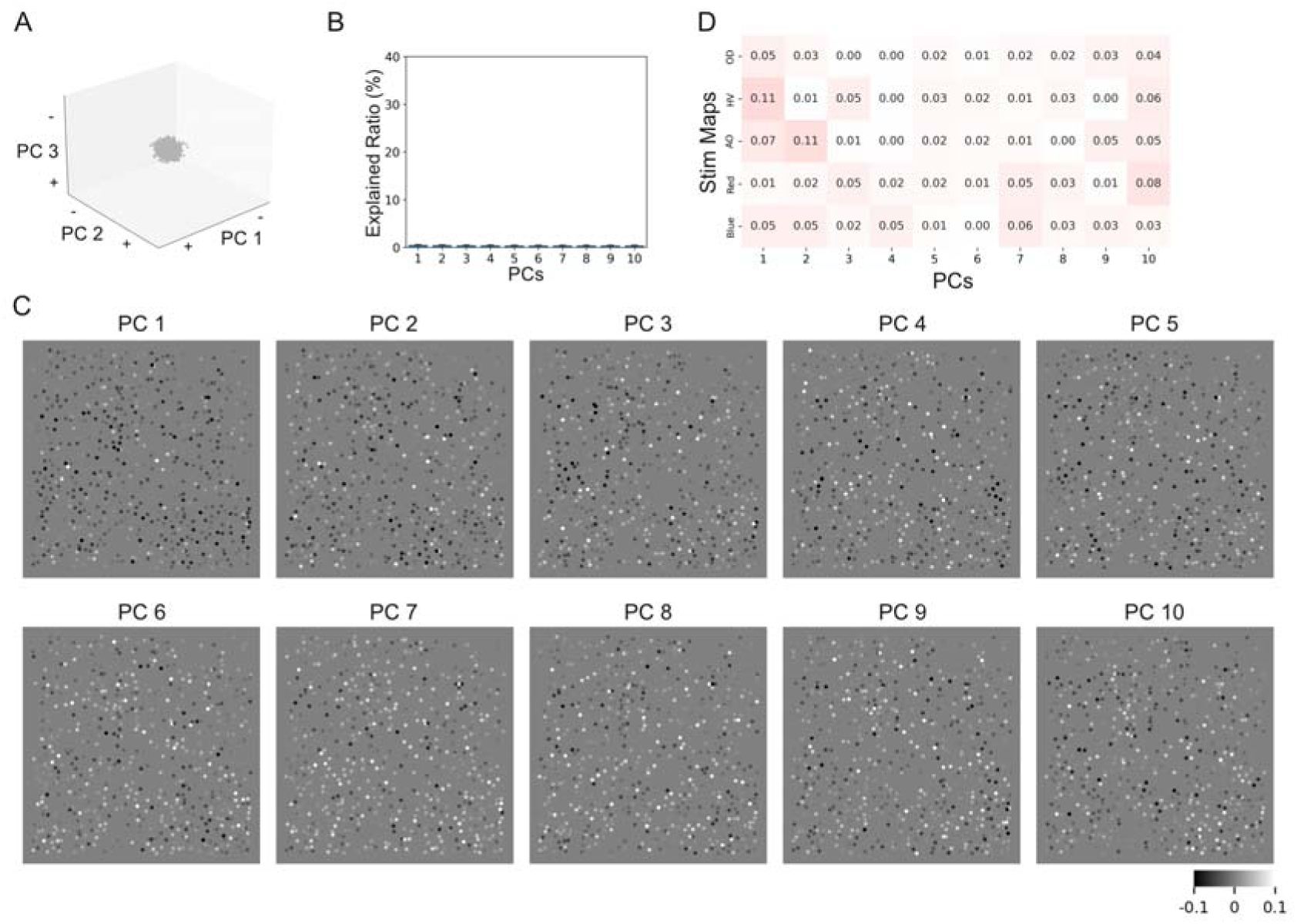
PCA on shuffled spontaneous data. **A-D**. Similar to Figure 2B-D, but for shuffled data. Percentages of data variance accounted for by the top 10 PCs were very low (sum of the top 10: 4.1±0.7%). There were no meaningful patterns in the PC visualization images.

**Figure S5.**
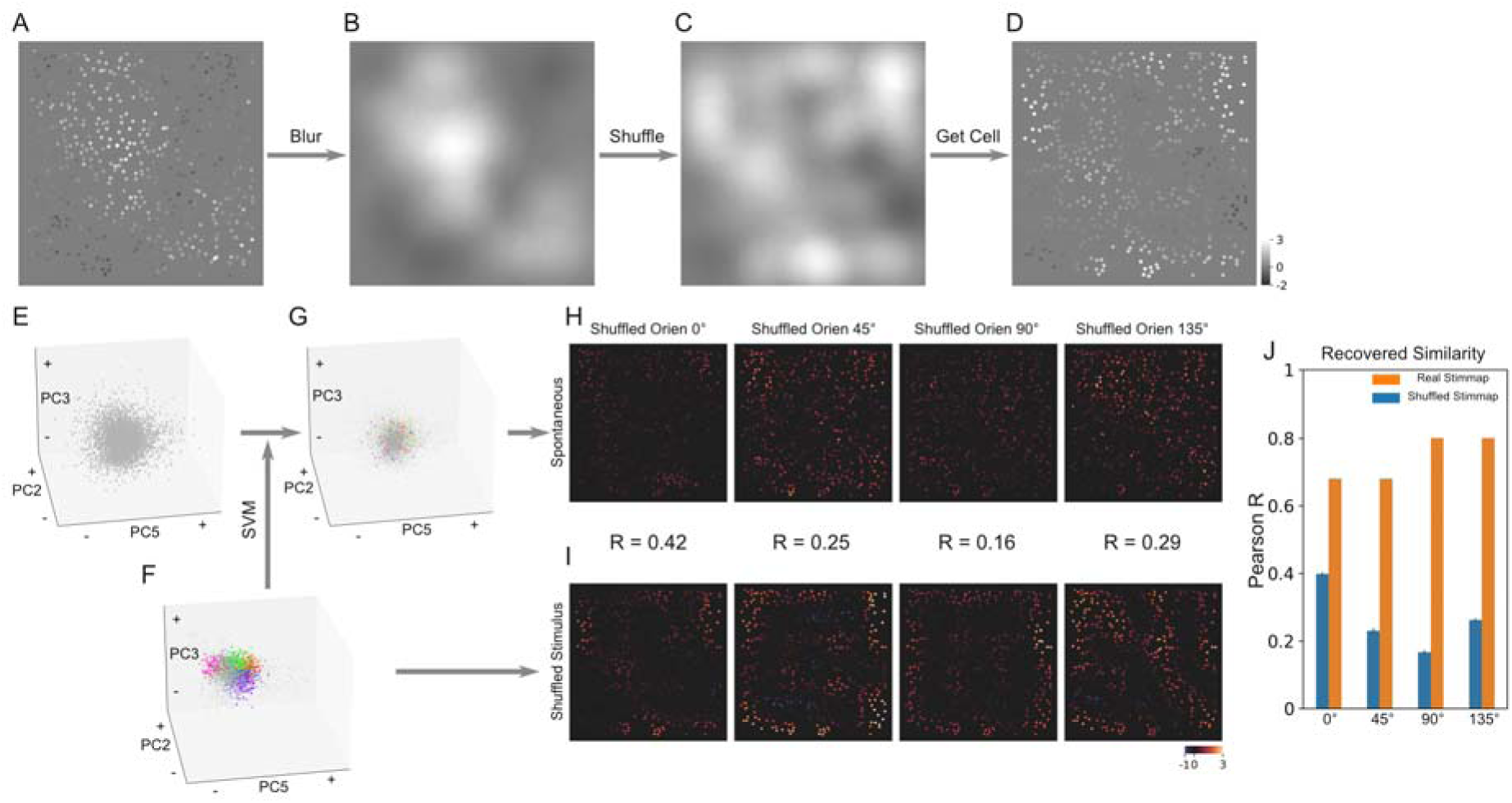
Negative controls for the SVM classification method. In classification of spontaneous data (Figure 3), we trained a SVM classifier with stimulus data (neurons’ response to different oriented gratings), and found stimulus-like patterns in the spontaneous data. It is not known whether an arbitrary pattern can be found in the spontaneous data if a SVM classifier is trained with such patterns. Theoretically, any pattern can be found in an infinitely long spontaneous imaging. Here we trained a SVM classifier with artificially generated patterns to provide a control for this method. **A-D**. Generation of artificial stimulus data based on data used in Figure 3. A real orientation-activation pattern for neurons (A) was first blurred (B) and phase shuffled to generate a new frame pattern (C). A new neuron-activation pattern was generated (D) based on this frame pattern. The same procedures were repeated for 8 different orientations. **E-I**: Same as in Figure 3A-E, but used artificial orientation frames for training. Orientation training frames (n=1451) were generated by adding random variations to neurons’ activation in the artificial frame in D. Training frames were plotted in the same PCA space (F) obtained after PCA analysis for spontaneous frames (E). Spontaneous frames can be classified as orientation frame (G) using the trained SVM classifier. However, the patterns of these classified spontaneous frames (H) did not resemble the artificial orientation patterns (I). **J**: Overall, correlation values from real classification were higher than those obtained with the artificial data.

**Figure S6.**
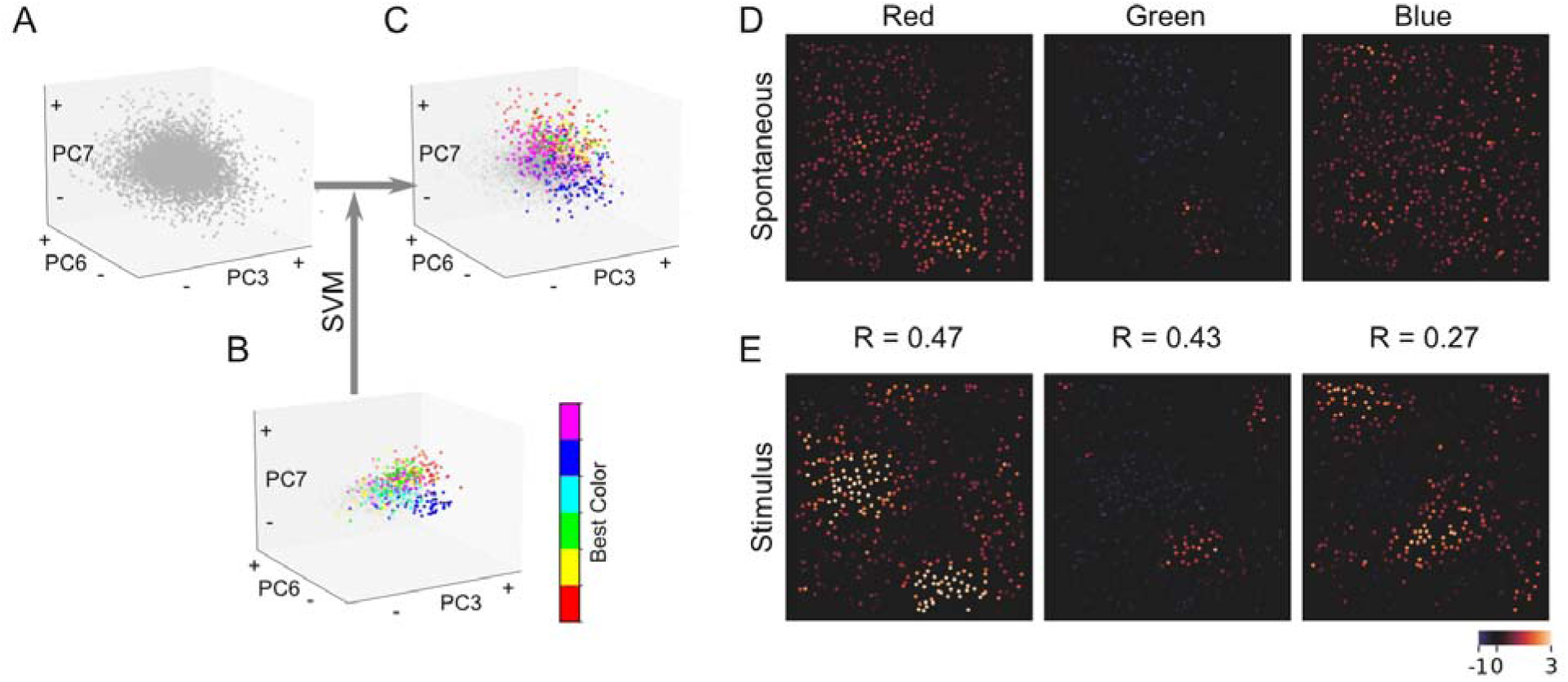
Spontaneous color patterns in V1. **A-E**. Similar to Figure 3 A-E, but for color classification. Although some frames were classified as color frames (B) and the averaged spontaneous color map had some similarity as the color functional maps (D&E). The overall correlations were low.

**Figure S7.**
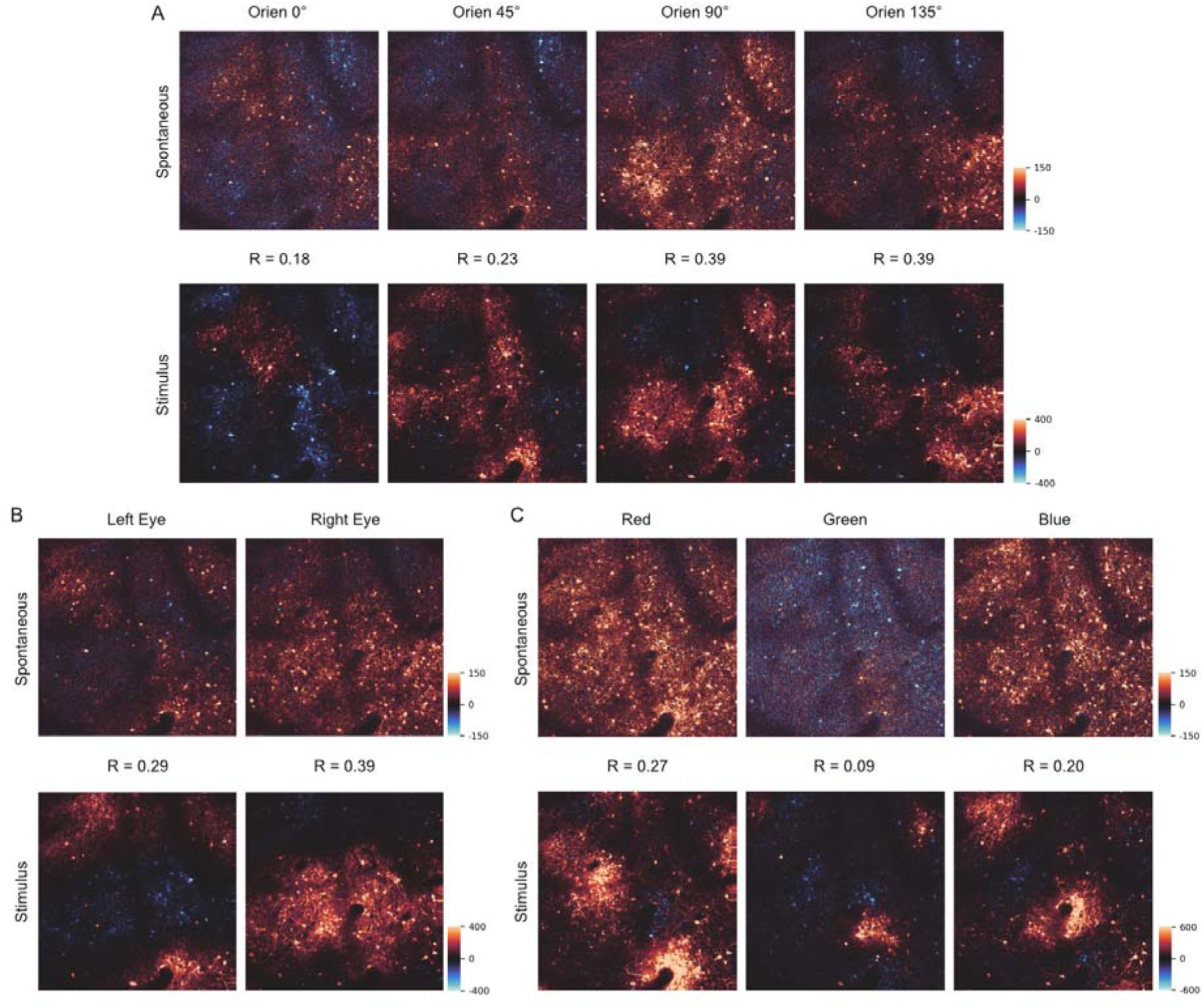
Pixel-based spontaneous patterns in V1. **A-C**: Similar to maps in Figure 3 D&E, I&J, and Figure S6 D&E, but calculated based on all pixels in the frames (averages of the dF frames, not the cell data). The similarity between spontaneous maps (top row) and functional maps (bottom row) was evident for orientation patterns (A) and ocular dominance patterns (B), but weaker for color patterns (right).

**Figure S8.**
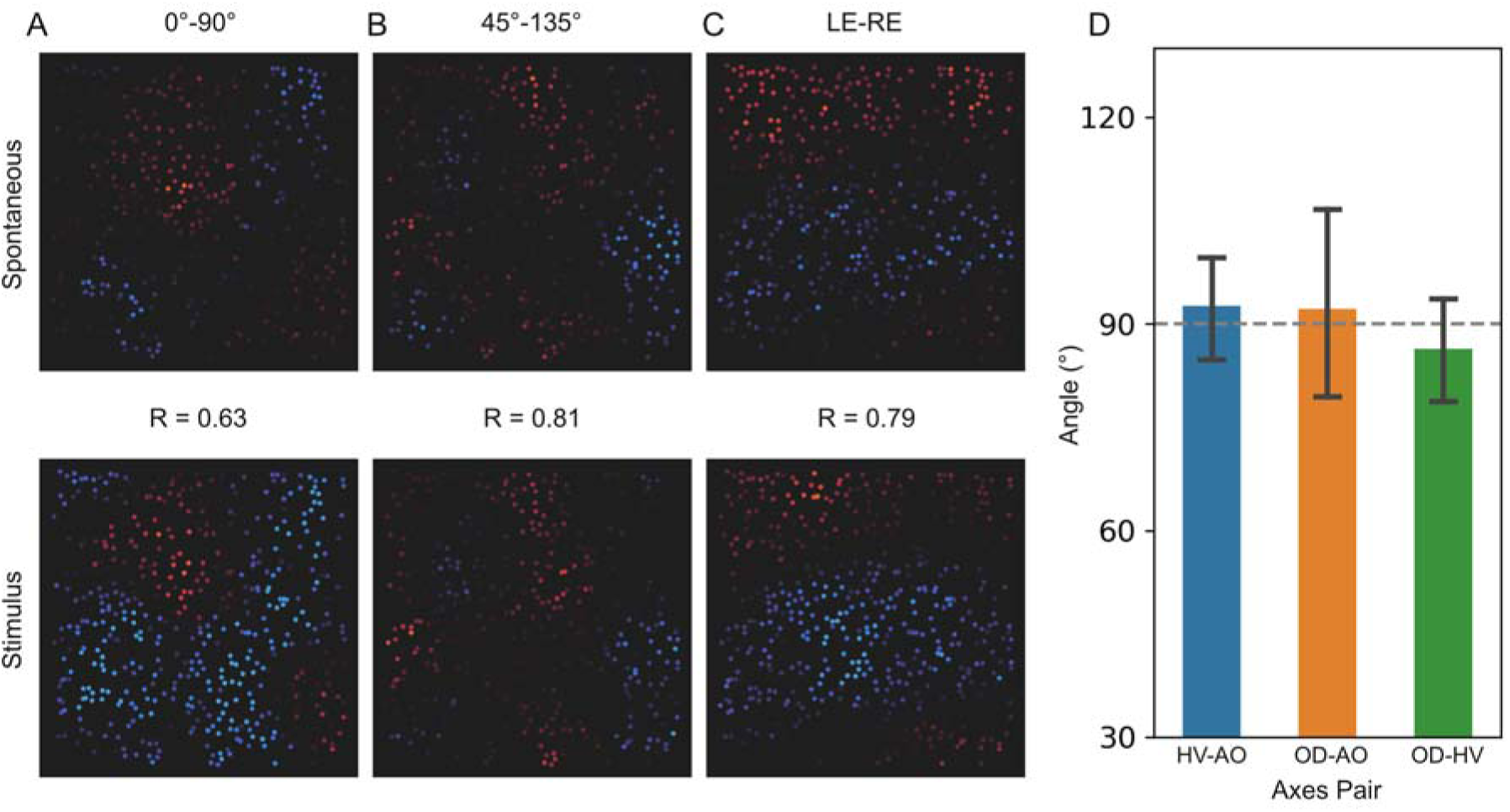
Spontaneous patterns from frame PCA. **A**. In the PC frame space shown in Figure 4C, 0°-preferring neurons and 90°-preferring neurons were spatially separated. We calculated the weighted sum of all the frame PCs using the mean 0° and 90° coordinates. Finally, we obtained a difference map by subtracting these two visualized maps (A). This 0° vs. 90° preference map had a good correlation with the 0° vs. 90° map obtained with stimulus-driven data, indicate that there was orientation information in the spontaneous data. **B**. Similar to A, but for 45° and 135° preferring neurons. The result map also had a good correlation with the 45° vs. 135° orientation map. **C**. Similar to B, but for LE and RE preferring neurons. It also had a good correlation with the LE vs. RE OD map. **D**. For the left-right eye difference vector and two orthogonal orientation vectors (arrows in Figure 4 C&D), we calculated their high-dimensional angles in the PC space. The three angles were all close to 90 degrees, indicating that these spontaneous patterns occurred independently, had very little temporal correlation.

